# Generation of transplantable and biologically responsive colonic tissue from human induced pluripotent stem cells using a rapid co-differentiation platform

**DOI:** 10.1101/2023.12.08.570795

**Authors:** William Dalleywater, Alexander V. Predeus, Batuhan Cakir, Pavel Mazin, Jayakumar Vadakekolathu, Sergio Rutella, Marian L. Meakin, Alison A. Ritchie, Shamir Montazid, Sara Cuevas Ocaña, Nadine Holmes, Victoria Wright, Fei Sang, Adam Bills, Declan Sculthorpe, Rasa Elmentaite, Sarah A. Teichmann, Shazia Irshad, Ian Tomlinson, Andrew Silver, Ricky D. Wildman, Nicholas R.F Hannan, Felicity R.A.J. Rose, Mohammad Ilyas

## Abstract

**Background:** The colonic mucosa consists of cell populations derived from multiple lineages. Induced pluripotent stem cells (iPSCs) are capable of generating large numbers of differentiated cells from any lineage. Thus, iPSCs are highly versatile for derivation of intestinal cells for generation of colonic mucosal tissue for clinical and biological applications.

**Objective:** We set out to create a human iPSC (hiPSC) multi-lineage co-differentiation platform capable of generating colonic mucosal tissue in vitro.

**Design:** We used hiPSCs and designed a differentiation protocol consisting of small molecules and recombinant growth factors to generate multiple cell lineages. Cells were seeded onto collagen hydrogels (forming colonic patches - CoPs) and modulated with multiple growth factors important in intestinal biology. CoPs were transplanted into immunosuppressed mice. Generated cells and tissues were profiled with transcriptomic analysis.

**Results:** hiPSC co-differentiation led to multiple intestinal epithelial, mesenchymal and endothelial cell populations. Seeded onto collagen scaffolds these cells created CoPs, which were transplanted into mouse subcutis. Engrafted CoPs developed into normal-looking colonic mucosa containing epithelial crypts (with enterocytes, goblet cells and neuroendocrine cells), multiple lamina propria-resident stromal populations and muscularis mucosae smooth muscle. They anastomosed to murine vasculature and maintained in-vitro for several weeks. We demonstrated that CoPs respond to known signalling pathways important in colonic mucosal biology and fibrogenesis, showing potential to provide a complex model of colonic pathobiology.

**Conclusion:** This platform could offer an accurate model of intestinal pathobiology, supply cells for regenerative cell therapies to treat intestinal disease, and provide therapeutic autologous grafts to repair damaged colon.

## Introduction

Diseases of the colon are common and constitute a major health burden. Transplanting functional, healthy intestinal tissue could be a novel therapeutic approach, particularly for chronic diseases such as inflammatory bowel disease (IBD). The colonic mucosa is the main functional layer of the colon and is thus a focus for colonic regeneration therapies(1,2). Loss of mucosa is a component of the pathological triad of IBD and thus restoring healthy mucosa may aid in breaking the cycle of chronic inflammation and microbial colonisation, to aid healing(2–4). There is also an urgent need for accurate experimental mucosal models to investigate disease mechanisms and identify novel targets for therapy.

The colonic mucosa is characterised by epithelial crypts supported by a range of mesenchymal cells including fibroblasts and endothelial cells and a deep band of smooth muscle, called *muscularis mucosae*(5–7). These cell populations derive from endoderm and splanchnic mesoderm embryologically and arise in tandem from the developing gut tube. Other cell populations, such as haemopoietic and neural cells migrate in during embryonic life from other germ layers(7).

Since colonic pathologies often involve complex interactions between epithelial and mesenchymal populations (containing multiple cell types(6)), any model should contain all these cell types. Investigating disease biology is further complicated as stroma is now known to contain several cell populations, which are precisely localised in the lamina propria (such as the peri-crypt fibroblasts and *SOX6* expressing cells around the stem cell niche(6,8)). Differing populations may therefore have different relationships with epithelium(3,6). Finally, since cells of the colon form a dynamic ecosystem, pathological change in one mesenchymal population may result in changes to other mesenchymal or epithelial cell populations.

The most popular existing intestinal models are *in-vitro* organoids and recent improvements in methodology have increased their utility. However, these models require access to primary tissue (usually surgical resection specimens) and are not able to replicate fully the complex cellular population profile and 3D colonic architecture(9,10). Mouse disease models are an alternative with certain advantages; but are expensive, can be difficult to manipulate and, critically, observations in mouse models may not translate to humans.

Our aim was to generate human intestinal cell populations from hiPSCs and illustrate their potential to reconstruct normal colonic mucosal tissue *in vitro* and *in vivo*. For successful *in vivo* engraftment, we reasoned that a transplant should resemble, as much as possible, normal colon in organisation and cell types. We hypothesised that a protocol with reproducible co-differentiation of hiPSCs into various cell populations present in normal colonic mucosa would represent a more developmentally relevant approach, given the importance of inter-lineage cues in shaping differentiated cell phenotypes(5,11–13). These would then need to be grown on an *in vitro* scaffold allowing assembly into architecturally normal mucosa, and which would be robust enough to allow transplantation. We reasoned that such a cellular construct should respond to known biological cues in colonic mucosa, therefore tested a selection of key growth factors involved in mucosal maintenance and disease.

Here, we present a rapid and robust co-differentiation protocol for generating intestinal cells from hiPSCs that can develop into normal colonic mucosa both *in vitro* and *in vivo* and is biologically responsive.

## Results

### Defining a novel serum-free protocol

To replicate the complexity of the mucosa, we aimed to co-differentiate hiPSCs simultaneously to form endoderm and mesoderm. We favoured a co-differentiation approach as this would allow interactions between different cell populations during differentiation, mimicking early developmental events in gut tube and early intestinal development. Many published protocols rely on poorly defined media components which are prone to batch-to-batch variation (such as conditioned media and animal derived serum) or use antibiotics(14–19). As our aim was to design a protocol that could be suitable for eventual transplantation and for easy adoption in other laboratories for use in *in vitro* models, we reasoned that using serum-containing components would limit application(15). Thus, we set out to explore protocol options in which differentiation could be achieved with recombinant or small molecules only. The resulting co-differentiation protocol replaced serum with defined products, reduced recombinant proteins and was conducted in antibiotic-free conditions. We tested this novel protocol against two alternatives that contain the major non-serum components utilised in other widely used published protocols(15,20).

### Novel protocol leads to co-differentiation of hiPSCs into endoderm and mesoderm

By day 4 of the novel protocol (protocol 1), there was up-regulation of endoderm markers *SOX17* and *FOXA2* and the mesoderm marker, *TBXT* (Brachyury) which was not seen in comparative protocols (Fig.1B)(20). All protocols showed a strong downregulation of pluripotency markers, *POU5F1* and *NANOG*.

**Figure 1:**
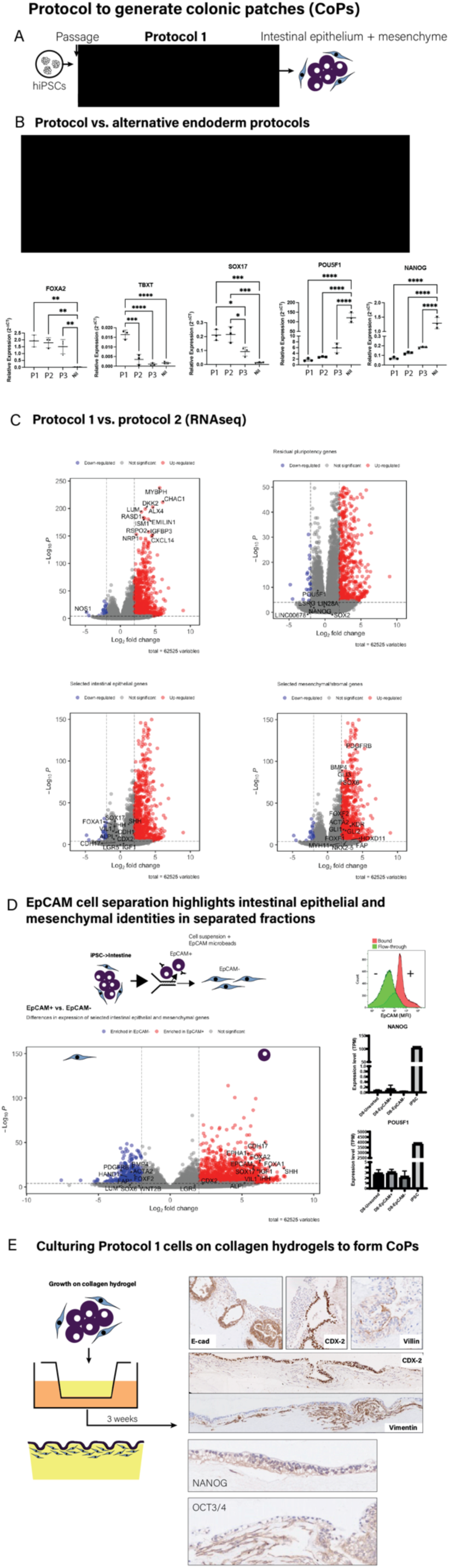
A novel protocol capable of deriving multiple intestinal cell populations. **(A)** Overview of the novel differentiation protocol **(B)** Comparison of protocol with alternative protocols. Expression of endoderm (*FOXA2, SOX17*)/mesoderm (*TBXT/Brachyury*) and pluripotency (*POU5F1/NANOG*) markers by qRT-PCR at day 4 of each protocol demonstrates enhanced co-differentiation with the novel protocol (see methods) (n=3 independent experiments). Error bars indicate standard deviation. Statistical significance assessed by one-way ANOVA. **(C)** The novel protocol results in enrichment of mesenchyme-related genes (e.g. *HAND1, LUM, PDGFRB, TNNT1*) (5, 6) compared with alternative protocol 2 (20). Cells were differentiated according to the protocols shown and then compared by mRNA-sequencing (n=6). Volcano plots illustrate top differentially expressed genes and specific epithelial and mesenchymal subsets. **(D)** Protocol 1 co-differentiated cells when MACS separated using EpCAM expression showed enrichment of intestinal epithelial genes in the EpCAM+ fraction (e.g. *CDH17, EPCAM, VIL1, ALPI*) and intestinal mesenchymal genes in the EpCAM-fraction (e.g. *ACTA2, BMP4, SOX6, LUM*). Cells were differentiated and then separated based on EpCAM expression (MACS) and profiled by flow cytometry/mRNA-sequencing. Representative flow cytometry plot following separation by MACS® cell separation of separated populations (n=3). Volcano plots demonstrate top genes enriched in EpCAM+ and EpCAM-populations. **(E)** Stratification into intestinal epithelium and mesenchymal self-assembled layers was shown when differentiated cells, as a mixed population, were seeded onto collagen hydrogels (2mg mL^-1^) and cultured *in vitro* for 3 weeks and assessed for E-cadherin (E-cad), CDX-2, Villin-1, Vimentin, Nanog and Oct3/4 expression by immunohistochemistry, representative of n=3 independent experiments.

**Figure S1:**
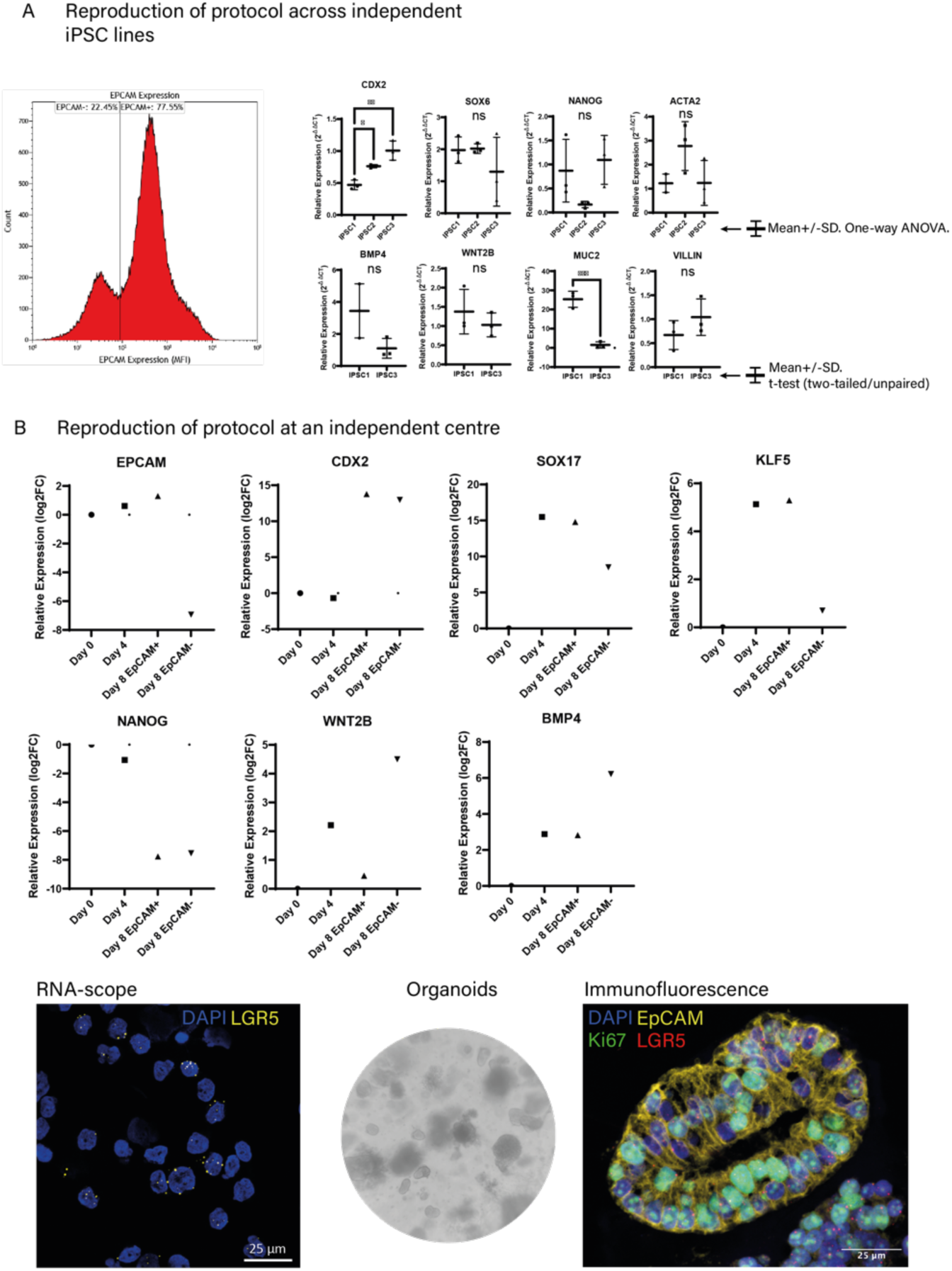
The differentiation protocol shows consistency across different hiPSCs and laboratories. **(A)** The co-differentiation protocol was applied to three different hiPSC lines 1-3 derived from both males (REBL-PAT; n=1) and females (BE-31, BE-32; n=2) demonstrated consistency of intestinal epithelial and mesenchymal co-differentiation across multiple hiPSC lines. Differentiated populations consisted of both EpCAM+ and EpCAM-cells (representative flow cytometry histogram from one alternative iPSC line). Evidence of intestinal epithelial (*CDX2, VIL1, MUC2*) and mesenchymal co-differentiation (*ACTA2, WNT2B, SOX6, BMP4*) was demonstrated by qRT-PCR (n=3, one-way ANOVA) as well as suppression of *NANOG* (comparison with hiPSC3 – REBLPAT hiPSCs). Central bar indicates mean and error bars indicate standard deviation. Ns = not significant. **(B)** The co-differentiation protocol was applied to a fourth hiPSC line (OSTI GI4380) by collaborators at an independent research facility (SM, SI, IT) and showed evidence of intestinal epithelial and mesenchymal co-differentiation. Evidence of intestinal epithelial (*EPCAM, CDX2, SOX17, KLF5*) and mesenchymal (*WNT2B, BMP4*) co-differentiation was shown as well as suppression of pluripotency (*NANOG*) (n=1). EpCAM+ cells were evaluated for human *LGR5* mRNA by RNA-scope (25% across 3 independent replicates) and could form organoids with a mixture of cell types (EpCAM – yellow, Ki67 – green, LGR5 – red, DAPI – blue), representative of n=6. After MACS separation, EpCAM^+^ cells were fixed on a coated slide using cytospin and stained with RNAScope probe for Human *LGR5* mRNA. 7500 cells from 3 independent replicates were counted; 25% of EpCAM+ were positive for *LGR5*, while no cells were positive for LGR5 at day 0

We identified transcriptomic profiles by RNA-Seq at Day 8 in cells cultured under protocol 1 and undertook a comparison with cells cultured using alternative protocol 2 (Fig.1C, Table 1)(15,16). Differentiated cells derived using both protocols showed similar levels of expression of intestinal epithelial markers (Fig.1C, lower left panel), but cells grown under the novel protocol were also strongly enriched for stroma-related genes including *HAND1, LUM, BMP4, ACTA2, SOX6* and *KDR* (Fig.1C, lower right panel)(5,6).

We further validated cell identity by separating cultures at the end of the novel protocol into enriched “epithelial” and “mesenchymal” groups using magnetic cell separation (MACS) based on EpCAM expression (a marker of epithelium) (Fig.1D). Expression profiling by mRNA-Seq confirmed intestinal epithelial identity of the EpCAM+ group (compared with the EpCAM-group) including enrichment of general epithelial markers (*SOX17,LGR5,SHH,IHH)* and specific intestinal markers (*CDH17,ALPI,VIL1*). Compared with the EpCAM+ group, the EpCAM-group showed enrichment for mesenchymal markers including *ACTA2, BMP4, LUM* and *SOX6* (5,6)(Fig.1D). There was no difference in pluripotency marker expression between the separated groups (Fig.1D). Moreover, expression was >1000 fold lower in differentiated cells compared with undifferentiated iPSCs.

From Day 9 onwards, co-differentiated cells (without MACS separation) were placed on collagen I hydrogels formed in Transwells to create “colonic patches” (CoPs) (Fig.1E)(21). These colonic patches were intended to simulate the organisation of the colonic mucosa, with exposure of the culture surface to culture medium. By Day 22 of *in vitro* culture the epithelial cells in the CoPs lined the surface of the hydrogel and had assembled into crypt-like structures. These cells expressed the intestinal epithelial markers, E-cadherin and CDX-2, together with Villin-1 (which demonstrated notable polarised expression limited to the apical cell membrane). Vimentin-positive mesenchymal cells in CoPs were stratified beneath the surface epithelium and around the crypts. No evidence of residual pluripotent cells was demonstrated by immunohistochemistry for OCT3/4 and NANOG. Thus, the colonic patches self-assembled into colonic mucosa-like structures with a polarised epithelium supported by an underlying mesenchyme.

Finally, our data were replicated and were consistent among four different hiPSC lines (Fig.S1A/B) including being performed independently by collaborators in another academic institution (Fig.S1B), thus demonstrating the robustness and the ease of implementing the protocol. In addition, EpCAM+ cells showed abundant *LGR5* transcripts by RNAscope, and formed proliferating (Ki67+) organoids when encapsulated in Matrigel.

### Determining effects of intestinal cytokines on mucosal CoPs

Colonic Patches (CoPs), as developed here, contain multiple mesenchymal and epithelial cell populations. We investigated the behaviour of the cell populations by stimulating established CoPs with specific cytokines (Fig.2A-C) to determine how well derived cell populations recapitulated known features of intestinal biology. In response to sustained CHIR-99021 stimulation (Fig.2A), epithelial and stromal compartments showed marked proliferation with cell stratification (similar to adenomatous tissue) seen in the epithelium. Treatment with either BMP4 or the Wnt inhibitor, ICG-001, caused a reduction in cell number. ICG-001-treated cultures showed extensive apoptosis demonstrating the requirement of Wnt activity for cell viability in both the epithelium and the stroma. Interestingly, the baseline condition (no exogenous Wnt stimulation) showed sustained viability, suggesting that endogenous stromal Wnt ligands produced in CoPs are capable of meeting epithelial Wnt requirements. When treated with the fibrogenic cytokine TGF-β1 (Fig.2B), the mesenchymal compartment of CoPs took on a well-described myofibroblast molecular profile characterised by expression of genes related to matrix-synthesis and contractile-proteins (*COL1A1*^high^/*ACTA2*^high^). *TIMP1* was upregulated but there was no change in *MMP1* expression. These changes were all abrogated if the CoPs were treated with A83-01 (a potent inhibitor of TGF-β receptors, ALK4, 5 and 7)(3,5,6). TGF-β1 stimulation was accompanied by contraction of the CoP, features resembling fibrous scarring seen in IBD, and deposition of extracellular matrix was confirmed with picrosirius red stain (Fig.2B). Treatment of CoPs with A83-01 (without TGF-β1 stimulation) prevented remodelling and fibrosis of the CoPs demonstrating intrinsic TGFβ1 activity in CoPs and further supporting CoPs as a model of normal colon.

**Figure 2:**
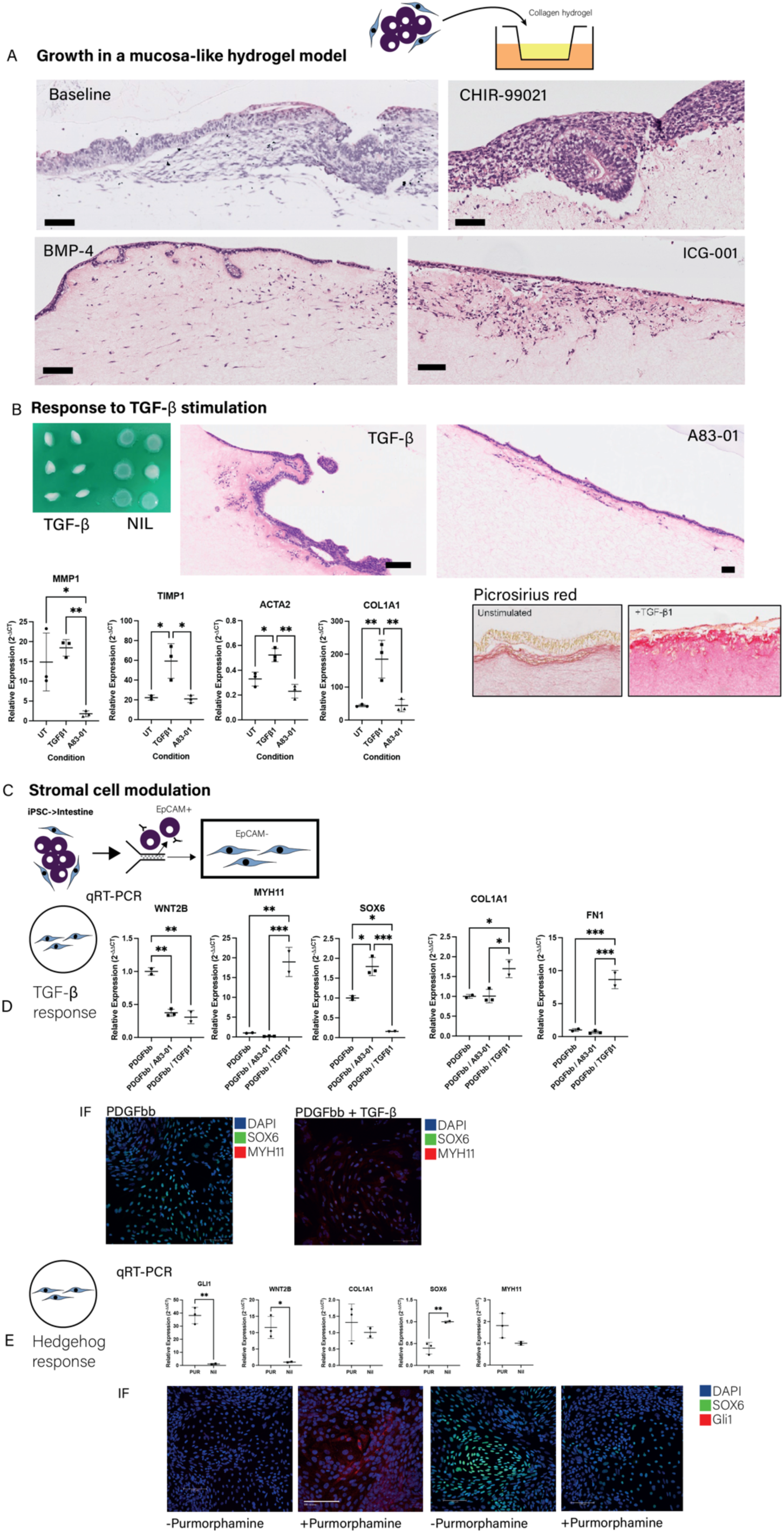
iPSC-derived CoPs respond to colon-related growth factor cues to model colon mucosal biology. (A) hiPSC-derived intestinal cells were seeded onto collagen hydrogels and then cultured for three weeks in the presence of CHIR-99021 (5 μM), BMP-4 (5 ng/mL) or ICG-001 (1 μM). Representative H+E stained photomicrographs of at least two independent replicates. Scale bars = 100 microns. (B) TGF-β induced a fibrosis-like response in CoPs. hiPSC-derived intestinal cells were seeded onto collagen hydrogels and then cultured for three weeks in the presence of TGF-β1 (1 ng/ML), A83-01 (500 nM). Representative H+E or PSR stained photomicrographs of at least two independent replicates. Scale bars = 100 microns. Gene expression was evaluated by q-RTPCR (n=3, one-way ANOVA). Central bar indicates mean, error bars indicate standard deviation. (C) Schematic showing intestinal stromal cells that were first differentiated from hiPSCs and then grown independently in 2D culture by depleting epithelial cells using MACS EpCAM separation. (D) TGF-β1 induced a myofibroblast-like phenotype in EpCAM-hiPSC-derived intestinal stromal cells. Separated EpCAM-cells were treated with PDGFbb (2ng/mL) and TGF-β1 (1ng/mL) or A83-01 (500nM) for 1 week and then assessed for gene expression (*SOX6, WNT2B, MYH11, FN1, COL1A1*) by q-RTPCR (n=2/3, one-way ANOVA). Central bar indicates mean, error bars indicate standard deviation (SD). Expression of SOX6 (green) and MYH11 (red) proteins was assessed by immunofluorescence with DAPI nuclear counterstain (blue) (micrographs representative of n=3). (E) Purmorphine induced a crypt-base fibroblast-like phenotype in EpCAM-hiPSC-derived intestinal stromal cells. Separated EpCAM-cells were treated with or without purmorphamine (10 μM) for 1 week and then assessed for gene expression (*GLI1, COL1A1, SOX6, MYH11, WNT2B*) by q-RTPCR (n=2/3, unpaired t-test). Expression of SOX6 (green) and Gli1 (red) was assessed by immunofluorescence with DAPI nuclear counterstain (blue) (micrographs representative of n=3).

To determine whether stromal cells could be further differentiated to recently described mucosal stromal subsets(5,6,22), EpCAM-cells were cultured in isolation and stimulated with different growth factor combinations (Fig.2C-E). When a monolayer culture of stromal cells was treated with only TGFβ1 supplemented in the media, cell proliferation was inhibited and we found that cell number and RNA quantity was insufficient for analysis. It was therefore necessary to add mesenchymal mitogen PDGFbb to increase cell numbers for the experimental work. We stimulated cells with TGFβ1 and A83-01 and investigated several genes related to the myofibroblast profile (*MYH11, COL1A1, FN1*) and pericryptal fibroblast profile (*SOX6^high^*)(6). We showed that on treatment with TGF-β1 (+PDGFbb), stromal cells took on a myofibroblast profile (*COL1A1^high^, FN1^high^, MYH11^high^, WNT2B^low^, SOX6^low^*) (Fig.2D). Stimulation with A83-01 (in combination with PDGFbb, but without TGF-β1) showed *SOX6^high^*, *WNT2B^low^*, *MYH11^low^*, *FN1^low^*-with no change in *COL1A1*. At the protein level, addition of TGF-β1 suppressed SOX-6 expression while Myosin-11 was only detected under TGF-β1 treatment conditions. These data show that, in the absence of epithelial cells, TGF-β1 appears to inhibit the pericryptal fibroblast profile whilst promoting the myofibroblast profile. More importantly, they show the flexibility and utility of our model.

We sought to mimic a population of pericryptal fibroblasts described previously as a characteristic of Hedgehog-responsive stromal cells surrounding the colonic crypt niche in mice(8) by treating stromal cells with purmorphamine (Fig.2E). We found an increase in *GLI1* and *WNT2B*, while SOX6 was downregulated, and myofibroblast-related genes were unchanged. This was supported at the protein level by a reduction in SOX-6 and increase in GLI1 after purmorphamine treatment. In previous studies, *SOX6* expression has been described as associated with BMP ligand production (i.e. a crypt apex localisation)(6). The inverse relationship of *GLI1/WNT2B* and *SOX6* expression is thus unsurprising and suggests a similar phenotype to that observed in Gli1+ crypt-base mesenchyme in mice.

### Single-cell RNA sequencing confirms co-differentiation of hiPSCs into multiple epithelial and mesenchymal sub-populations

To characterise the cellular populations in the epithelial and mesenchymal compartments, sc-RNAseq was performed following cell sorting (EpCAM+ and EpCAM-cells) immediately at the end of the novel protocol on day 8 and following a further 6 days of *in vitro* monoculture of EpCAM-cells (Fig.3A/B). From day 8 to day 14, EpCAM-cells were cultured with or without purmorphamine to study differentiation of the stromal population in more detail. Single-cell expression profiles of cells were compared to publicly available single-cell reference atlases generated as part of the human cell atlas project, using CellTypist (“Developing_Human_Organs”) (Fig.3C)(3,5,23). This demonstrated a diversity of emerging intestinal epithelial and stromal populations indicative of mesoderm differentiation towards the cell types seen in the mature intestinal stroma(23,24). The epithelial cell population was separated into subsets from the main dataset and mapped to the human gut cell atlas (“Cells_Intestinal_Tract”); revealing that epithelial cells mapped predominantly to “distal progenitors” as well as “transit amplifying” cells, supporting colonic identity (Fig.3D). The identity of all populations was further confirmed by demonstrating enrichment of marker genes for intestinal epithelium, endothelium, mesoderm and more mature mesenchyme (Fig.3E). Furthermore, we demonstrated a posterior homeobox programme (important for specifying the fate of the mesoderm towards an appropriate intestinal mesenchyme), dominated by *HOXA10/11, HOXB9* and *HOXC6-9* (Fig.3F). Our data showed expression of *CDX2* in both epithelial and mesenchymal populations at day 8, which are consistent with a recent finding that *CDX2,* although ultimately expressed only in mature epithelial tissue, is required for both intestinal endoderm and mesoderm fate specification(24) (Fig.3G). Within later mesenchymal clusters (clusters annotated 3+4, Fig.3B/G) subclusters of cells showed enrichment for expression of smooth muscle markers (*MYH11,ACTA2,CNN1,ACTG2*) whilst others demonstrated a phenotype typical of specific mature fibroblast subsets (*CXCL14,F3,RSPO2,NPY*)(5). The Hedgehog pathway is important for directing intestinal stromal lineages, as demonstrated in earlier experiments where stromal cells were stimulated with purmorphamine. We found that Hedgehog ligands were restricted to the epithelial population (Fig.3G). To understand further the effects on population profiles, stromal cells were profiled by single-cell RNAseq after treating with purmorphamine (Fig.3H). This caused enrichment of *GLI1* and *PTCH1* thereby confirming a Hedgehog response. Upregulation of *WNT4* and *SFRP1* and downregulation of BMP ligands were observed; features that recapitulate the phenotype of WNT-secreting crypt-niche mesenchyme described in mice(8).

**Figure 3:**
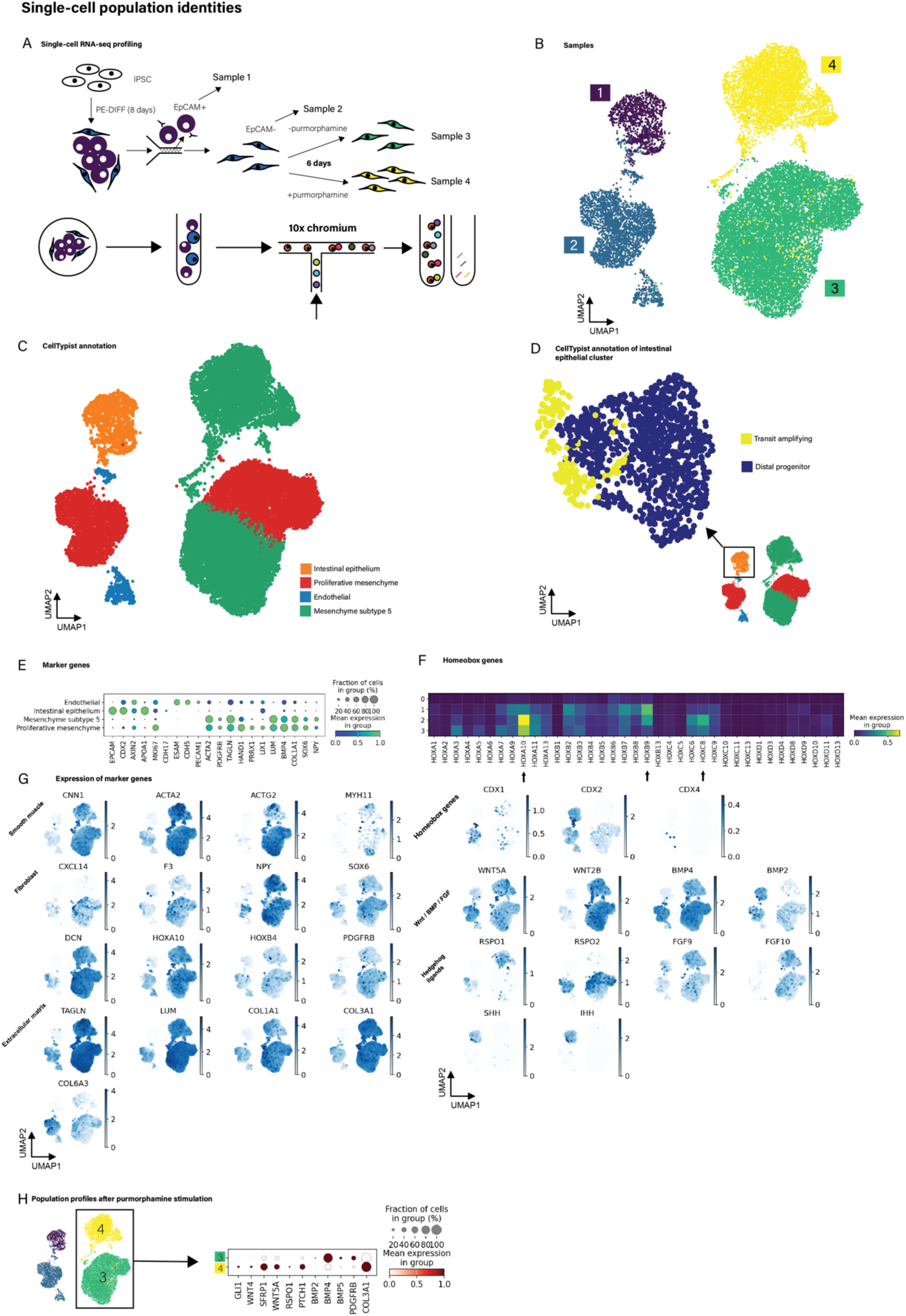
Single-cell RNAseq of separated cells after the novel protocol demonstrates intestinal epithelial, mesenchymal and endothelial population identities with expression of marker genes recapitulating intestinal population heterogeneity. **(A)** Overview of study design; sample 1: day 8, EpCAM+; sample 2: day 8, EpCAM-; sample 3: day 14 EpCAM-in monoculture after separation; sample 4: day 14 EpCAM-(monoculture) with purmorphamine treatment **(B)** Uniform manifold approximation and projection (UMAP) leiden clustering of all samples. **(C)** Automated annotation using the CellTypist algorithm of each cell population according to the “Developing Human Organs” model, showing intestinal epithelial, endothelial and mesenchymal identities. **(D)** Automated annotation using the CellTypist algorithm of the intestinal epithelial population according to the “Cells Intestinal Tract” model, showing distal progenitor and transit amplifying identities. **(E)** Dotplot of top marker genes of each population grouped by CellTypist (Developing Human Organs) annotations. **(F)** Mesenchyme (samples 1, 2, 3) are enriched for posterior homeobox genes. Heatmap showing expression of Hox genes in each sample, arrows indicate posterior Hox genes. **(G)** Feature plots of marker genes corresponding to smooth muscle and fibroblast lineages, Wnt/BMP/FGF and Hedgehog signalling pathways and extracellular matrix. **(H)** Comparison of mesenchyme with (sample 4) and without (sample 3) purmorphamine treatment to generate GLI1^+^/crypt niche phenotype, with a dotplot of key signalling and marker genes in each of the treated (sample 4) and control (sample 3) samples.

### From pluripotent stem cell to transplantable colonic patches (CoPs) in fifteen days without residual pluripotency

Given the wide array of cellular populations seen at Day 8 in CoPs with the novel protocol, we wanted to determine whether CoPs – even at this early stage – were suitable for transplantation and were capable of maturation to adult tissue. CoPs were modified by chemically cross-linking the lower part of the collagen gel (to increase strength for handling) while the upper part was not cross-linked (Fig.4A). Day 15 CoPs (eight-days of novel protocol and seven days of culture of the mixed cell population on collagen I) (Fig.4B/C) were transplanted into the sub-cutis of immunocompromised (*Rag2^-/-^ IL2RG^-/-^*) mice and harvested after 2, 3 or 4 weeks (Fig.4C/D). Across all experiments we found viable tissues consistently in around 55% mice indicating high transplant efficiency. This was higher than previously reported engraftment rates of approximately 10% achieved in a similar mouse model using a different protocol, where multiple separately derived primary intestinal cell populations cultured on tissue scaffolds were transplanted subcutaneously(25).

**Figure 4:**
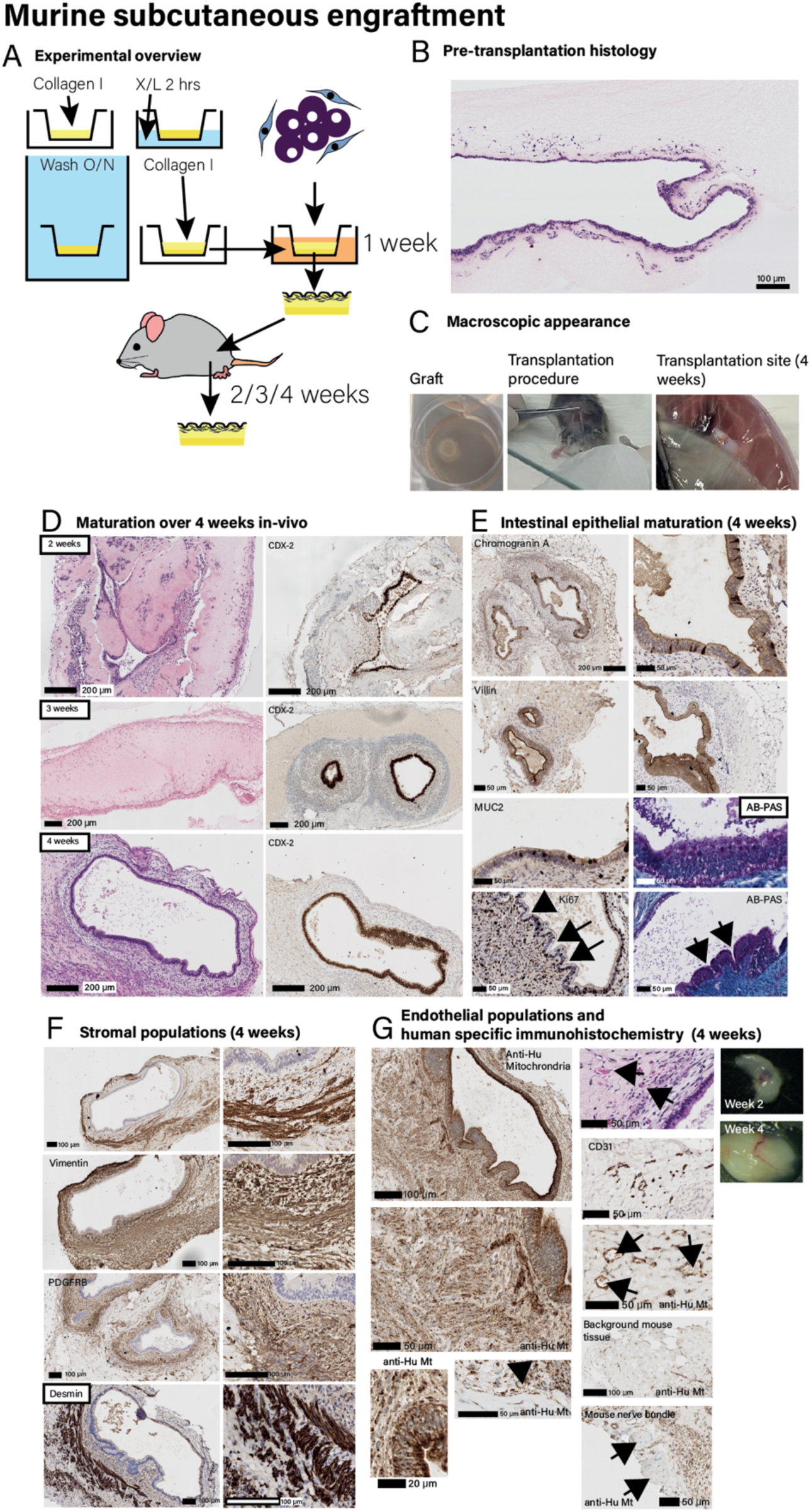
CoPs form mature stratified intestinal tissue within 4 weeks of murine subcutaneous engraftment. **(A)** Experimental overview of in vivo experiments A base layer of collagen I hydrogel was formed in 12 well plate Transwells from 2mg mL^-1^ neutralised collagen I and subsequently chemically cross-linked, after extensive washing, a further equal volume of collagen hydrogel (same concentration) was formed on top. Cells (mixed cell populations derived following the 8 day novel protocol) were then added and cultured for 1 week, before transplantation subcutaneously in immunosuppressed mice for 2 – 4 weeks. **(B)** Pre-implantation histology of CoPs shows stratification into epithelium on the surface with underlying mesenchymal cells. **(C)** Macroscopic appearance of transplant before engraftment (top), during transplantation (middle) and at sacrifice (bottom). **(D)** Images showing graft maturation over 2 – 4 weeks showing H+E stained tissue and CDX-2 immunohistochemistry. **(E)** Intestinal epithelial maturation at 4 weeks demonstrated by immunohistochemistry for Chromogranin A, Villin, MUC2 and Ki67 and AB-PAS stain for mucus. Arrows indicate formation of colonic crypts. **(F)** Stromal cell maturation and organisation at 4 weeks assessed by immunohistochemistry for smooth muscle actin, vimentin, PDGFRB and desmin. **(G)** Human origin of cell populations and functional human endothelium anastomosing with murine host shown by H+E stain, CD31 and human specific anti-human mitochondrial stain. Short arrows indicate vascular spaces of human origin, demonstrated by anti-human mitochondrial staining. Long arrows indicate mouse nerve bundle, with no staining by anti-human mitochondrial antibody.

**Figure S2:**
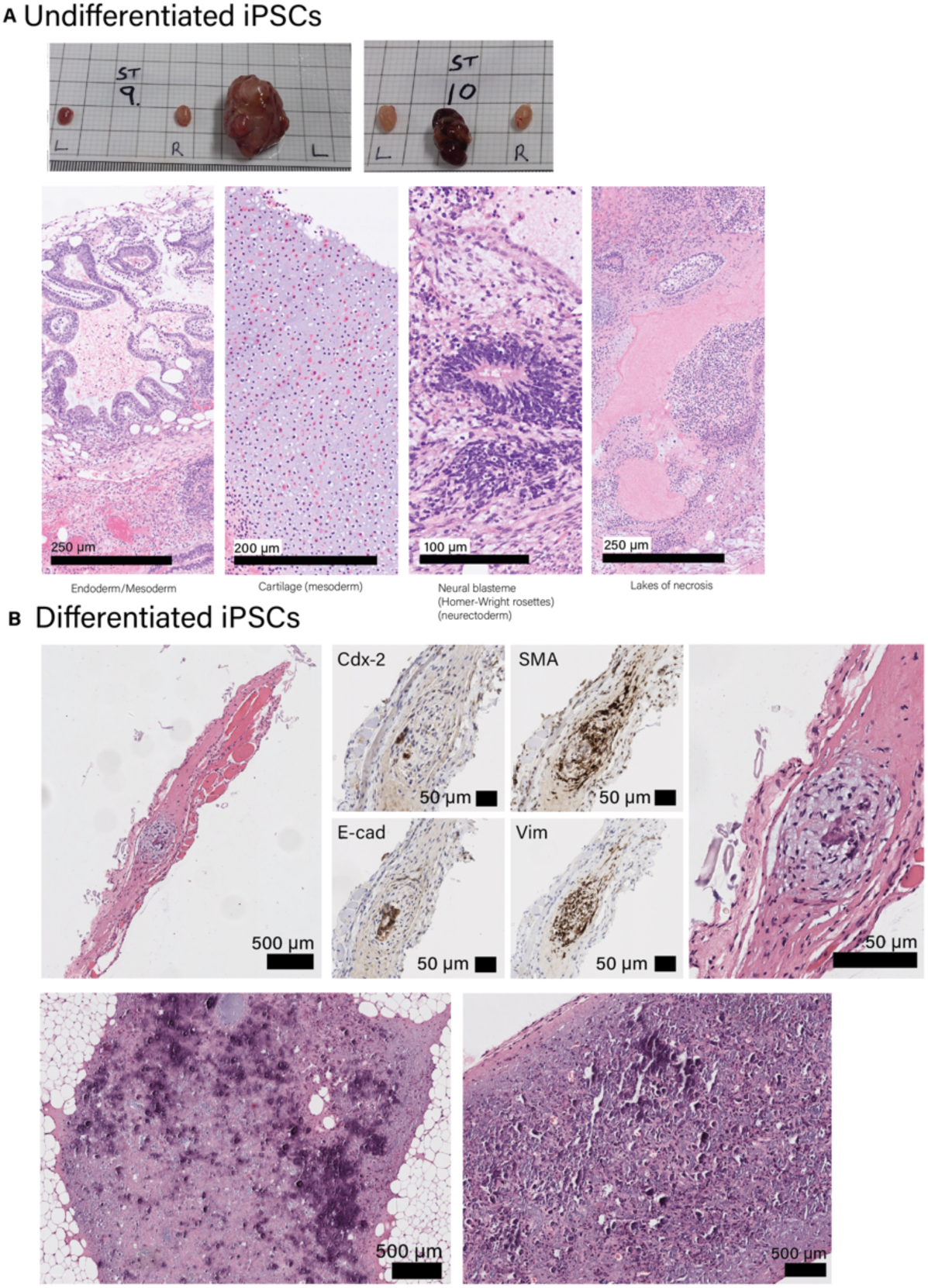
In vivo teratoma assay demonstrates no residual pluripotency in hiPSC-derived intestinal cells using the novel protocol. A) Undifferentiated hiPSCs (positive control): macroscopic photographs of tumours around the testicular subcapsule grown in immunosuppressed (Rag2^-/-^ IL2RG^-/-^) mice following implantation with undifferentiated hiPSCs. Representative photomicrographs (of twelve images) of H&E stained sections demonstrating features of tissues derived from all three germ layers. From left to right: endodermal / glandular tissue; cartilage (mesoderm); neural 24lastema / Homer-Wright rosette (ectoderm); lakes of necrosis. Scale bars represent distances indicated in individual photomicrographs. B) Differentiated hiPSCs (after novel protocol). All post-mortem tissues were examined by histology. Representative photomicrographs from each specimen where any possible engrafted tissues are shown. Tissue sections were stained with CDX-2, E-cadherin, SMA and vimentin of viable hiPSC-derived intestinal cells. Scale bars represent distances indicated in individual photomicrographs.

In our transplants, there was progressive increase in tissue organisation and cell maturation and, by four weeks, luminal spaces lined by epithelium with concentric layers of specialised mesenchyme emerged (Fig.4D). The epithelium showed intestinal differentiation including multiple cell types (neuroendocrine, goblet cells, and proliferative cells) (Fig.4E). Underlying mesenchyme resembled lamina propria and smooth muscle fibres (identified by Smooth Muscle Actin (SMA) and desmin expression) formed a boundary beneath this, recapitulating *muscularis mucosae* (Fig.4F). The tissues became highly vascularised and, using antibodies specific for human mitochondria, the vascular channels were shown to be lined with human endothelial cells (Fig.4G). The vascular channels contained blood indicating that human blood vessels had undergone anastomosis with murine dermal blood vessels to become functional.

One concern with transplantation of iPSC-derived tissues is the possibility of residual pluripotency leading to teratoma development. To confirm earlier molecular assays (Fig.1/S1) that the risk of teratoma had been removed through differentiation, we tested the cell populations in a teratoma assay performed at Day 8. This assay was performed in immunocompromised mice, which would permit any pluripotent cells to form teratomas. When undifferentiated hiPSCs were implanted into the mouse testicular subcapsular space (Fig.S2A), they formed large teratomas with all three germ layers represented. In contrast, when co-differentiated hiPSCs (Fig.S2B) were implanted, the transplants underwent degeneration with fibrosis and dystrophic calcification indicating apoptosis. Occasional intestinal glands were seen. No tumours formed and no other tissue type that may have derived from other germ layers was seen. These experiments indicate that differentiation fate specification is complete and there is no residual teratoma-forming activity following differentiation using the novel protocol. This model is relevant to human transplantation conditions where immunosuppression may be required for successful engraftment of tissue.

### Digital Spatial Profiling of transplanted colonic patches reveals maturation and development of gradients of normal biomarker expression along the crypt axis

The colonic mucosa is dynamic; epithelial cells migrate from crypt base to crypt apex over 3–5 days(26), differentiating from multipotent stem cells into terminally differentiated cells as they migrate(27). Tissue homeostasis is maintained by gradients of growth factors and cell surface receptors and these can be quantified using Digital Spatial RNA Profiling (DSP), allowing gene expression to be quantified in small regions of tissue.

Transplanted CoPs morphologically resembled colonic mucosa, thus we sought to show that this was the case transcriptionally by comparing DSP data from histologically normal human colon with CoPs sampled at 2 weeks, 3 weeks and 4 weeks after transplantation (Fig.5A). Endothelial transcripts were demonstrated in transplanted tissues (Fig.5B). Mesenchyme showed a progressive transcriptional shift over timepoints from an “immature” phenotype to a more mature phenotype (e.g.,decreasing *TWIST1*, mesoderm marker, Fig.5C). The mesenchyme retained *GLI1* expression, important for maintaining the stem cell niche, while epithelium became enriched for Hedgehog ligand, *IHH* (Fig.5D/E). Of particular note, was a gradient of increasing mesenchymal *PDGFRA* expression from the base of the lamina propria to the mucosal surface (Fig.5F) which coincided with epithelial maturation along the crypt axis. In normal adult colon tissue, we showed an opposing apex-base gradient of both *GLI1* (Fig.5E) and *PDGFRA* (Fig.5F) corroborating other studies which have demonstrated PDGFRA+ mesenchymal cells at the crypt apex(28). Our data suggest that the emergence of this population may be important for intestinal epithelial maturation, as demonstrated recently by Huycke et al.(29)

**Figure 5:**
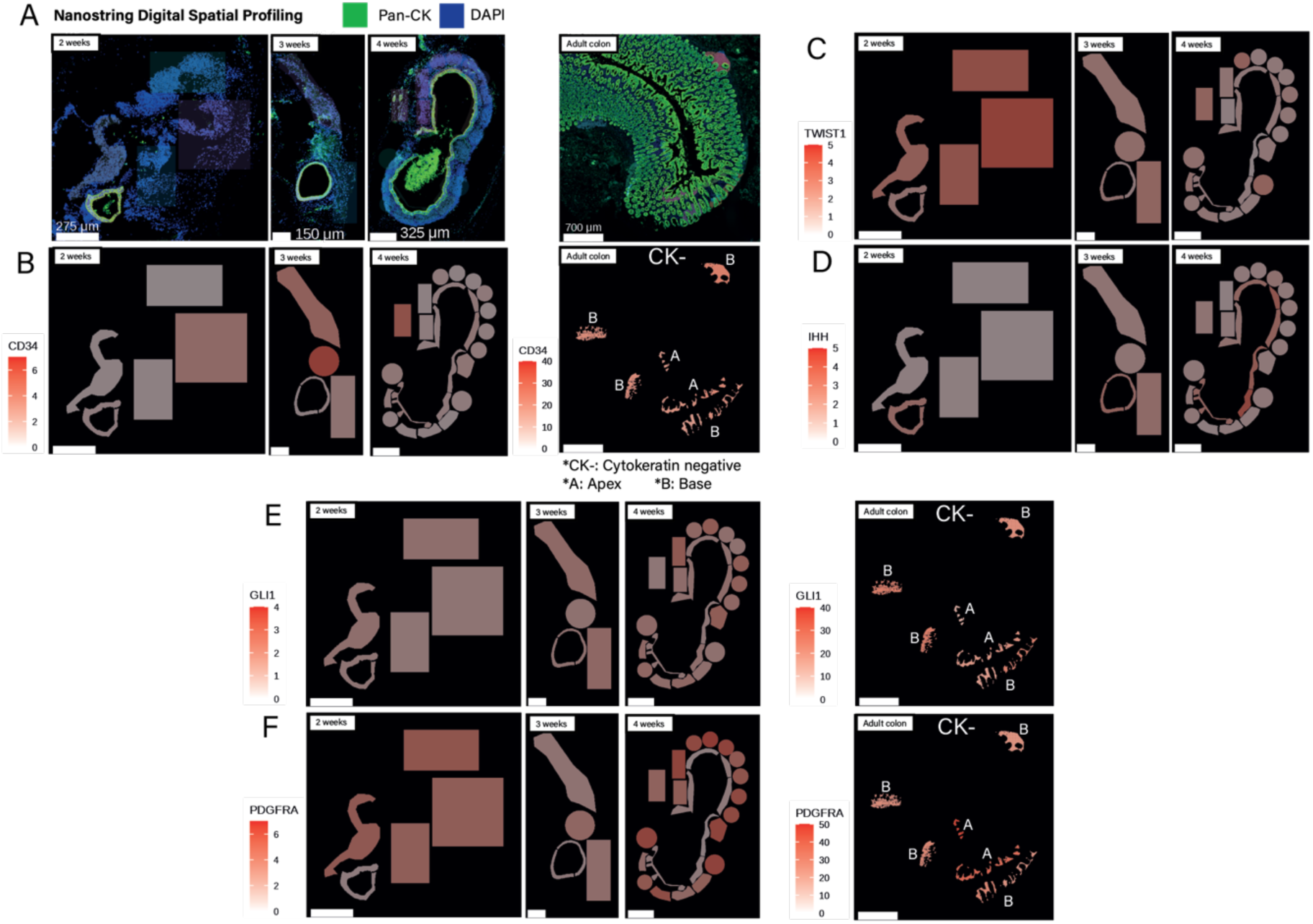
Nanostring Digital Spatial Profiling illustrating the maturation of cell populations within transplanted CoPs over several weeks of engraftment. (A) Pan-cytokeratin and DAPI stained iPSC-derived transplants at 2, 3 and 4 weeks and adult human colon control tissue (n=1 for each timepoint). All markers below correspond to the same tissues. (B-F) Demonstration of endothelial transcripts and maturation of mesenchyme during engraftment. (B) Endothelial cell marker, CD34, expression correlates with presence of vascular spaces in in vivo grafts and normal adult colon(Adult colon – CK-= cytokeratin negative cells, A: apex of crypt, B: base of crypt). (C) Mesoderm marker, TWIST1, expression decreases across 2 – 4 weeks timepoints of in vivo growth demonstrating maturation of mesenchyme. (D) IHH expression is limited to the developing intestinal epithelium (E) GLI1 (transducer of Hedgehog signalling) shows corresponding expression limited to the developing mesenchyme. GLI1 expression is higher in crypt base areas of illumination compared with crypt apices (F)PDGFRA shows increasing expression across 2 – 4 week timepoints of in vivo growth and is enriched crypt apex areas compared with crypt bases, suggesting mesenchymal PDGFRA expression may be required to drive more differentiated epithelial cell-types to emerge from in vivo hiPSC intestinal tissue grafts.

## Discussion

The availability of hiPSCs raises the possibility of engineering therapeutic autologous cell transplants to replace damaged tissue, or as *in vitro* tissue models. hiPSCs undergo differentiation into mature tissue derived from any or all the embryonic germ layers. To use hiPSCs as a cell source for the generation of healthy tissue, differentiation must be precisely controlled so that only the target tissue is produced. Furthermore, since a tissue rarely consists of only one cell type, all cell populations should be present and in the correct spatial location if the tissue is to be functional.

Here we have described a simple co-differentiation platform that can generate transplantable colonic grafts from hiPSCs within just 15 days. Unlike many published protocols, the novel protocol is serum-and antibiotic-free, facilitating translation into a clinically useful therapeutic or for use as a reproducible *in vitro* model. It has been replicated in several iPSC lines and in different laboratories illustrating the robustness of the protocol. The novel protocol generates multiple intestinal cell lineages simultaneously; confirmed by single-cell mRNA sequencing as well as *in vivo* engraftment studies. Co-differentiated cell populations do not contain residual pluripotency and they have been grown on biocompatible collagen gels (already in routine use in surgical applications) to create CoPs.

Finding cell sources for replacement therapies has largely focussed on derivation from native tissues or stem cell sources from adult tissue. Intestinal stem cells isolated from adult tissue can be cultured *in vitro* as 3D organoids that mimic the function and structure of the original tissue. These 3D stem cell units are self-organising and self-renewable, and usually cultured in Matrigel containing signalling factors important for the maintenance and regulation of the cells within the organoid(30). Organoid cultures of intestinal stem cells have played an integral role in our understanding of intestinal stem cell biology and its regenerative potential(10) and the concept of organoid regenerative medicine has been explored previously in animal models of IBD. Related studies have demonstrated that replacement mucosal cells can integrate into mucosa and aid healing in the disease tissue (1) illustrating the potential of a cellular regenerative medicine therapy to support healing in IBD. To consider organoids as a cell therapy, organoids would have to be fully characterised requiring extensive optimisation, be able to undergo mass expansion in a controlled, time-efficient manner, and their efficacy in human disease would need to be demonstrated(2). A further issue is that in diseased tissues it may not be possible to isolate healthy cells to then expand in cell culture. iPSC-derived intestinal cell generation offers several advantages over a primary intestinal organoid approach, particularly that protocols can rapidly generate large numbers of cells and multiple cell lineages can be generated simultaneously. Other iPSC studies have demonstrated some degree of co-differentiation but have considerably longer and more complex protocols including the use of serum, which we consider would limit clinical application(19). Compared with other published protocols, our method achieves rapid differentiation, with a greater diversity of cell types which form mature intestine. This method is simpler than others published in the literature (14,15,31,32) and makes use of defined media supplements which have the potential for either direct substitution with a GMP-compliant equivalent or will require minimal effort in refinement towards a GMP-compliant method.

To demonstrate maturation potential, xenograft transplantation into immunosuppressed mice is regarded as a gold-standard method for both intestinal organoid and iPSC-derived intestinal cells (16,25). Our subcutaneous model provides comparative data with other hiPSC-derived intestinal cells and primary intestinal organoid methods in which xenografts have been made at extra-intestinal sites including subcutis and renal. While in this study, we did not transplant xenografts into the intestine, the ability of our hiPSC-derived tissues to mature into colonic tissues outside the intestine is additional testament to their committed colonic differentiation without tissue-specific microenvironment cues. Transplanted CoPs into the *subcutis* of mice showed development of colonic mucosa complete with human-derived *muscularis mucosae* and human-derived blood vessels that successfully anastomosed to murine blood vessels. Compared with previous iPSC approaches(16,33), we saw similar levels of colonic maturation including ability to form smooth muscle bundles and anastomose with host vasculature, but with a shorter timeframe to transplantation from starting intestinal differentiation to transplantation (15 days vs. 37 days). We were able to be grow and mature our multi-lineage cells on the surface of hydrogel scaffolds to form CoPs, which we consider is likely to be more amenable to colonic transplantation in future. This scaffold model is similar to that developed by Meran et al.(25), where the authors were able to successfully transplant *in vitro* expanded primary intestine-derived jejunal tissue (rather than colon) into the subcutis of mice. We consider several advantages of an iPSC approach in comparison as (i) iPSC-derived cells could be grown on a relatively simple hydrogel platform and (ii) give rise to multiple populations of cells in a single protocol, compared with harvesting multiple cell populations. Our hiPSC-derived stromal populations replicate the full complexity of the intestinal stroma, as demonstrated by scRNA-seq studies(5,6). Thus, it follows that this may mediate appropriate epithelial crypt organisation, through Wnt and BMP signals, as well as releasing pro-angiogenic signals to promote vascularisation. This is exemplified by increased expression of PDGFRA over several weeks of transplantation, correlated to gradual acquisition of PDGFRA+ cells, which are known to drive intestinal epithelial differentiation(29). Finally, we anticipate the use of hiPSCs using our novel differentiation protocol offers greater opportunity for scale-up manufacture of these cell populations, unlocking the cell supply chain to produce tissue grafts for engraftment to the bowel to support healing.

We believe that CoPs represent a significant modelling development as they are the closest to normal colonic mucosa of any current *in vitro* model. The fact that our model responded in the expected way to stimulation with a variety of cytokines demonstrates validity of the model, such as in investigating progression of intestinal fibrosis development from normal mucosa and to test novel therapeutics to inhibit or even reverse fibrosis. Furthermore, hiPSC-derived CoPs grow on the surface of hydrogel scaffolds, giving rise to application for barrier / absorption studies. Another advantage of our model lies in the heterogeneity and responsiveness of stromal populations which would allow the study of specific diseases. For example, the SOX6+ population (also known in published literature as stromal 2 (6)) identified in our model is considered important in maintaining signalling pathways in normal intestinal crypts. Depletion of this population has been demonstrated in IBD(4). We showed that stimulation with cytokines active in IBD reduced numbers of SOX6+ cells and consequently reduced *SOX6* mRNA, indicating the potential of this model system for the study of IBD.

In summary, we have developed a simple colonic co-differentiation platform that derives multi-lineage cell populations from hiPSCs which can then be used to form transplantable colonic patches (CoPs) in 15 days. This represents a significant step in the development of a cell supply chain for regenerative cell transplants to treat colonic disease. CoPs also offer great potential for use as an accurate, dynamic and biologically responsive *in vitro* model for investigation of intestinal pathobiology.

## Acknowledgements

We are grateful for the support of all members of the research groups led by Profs Ilyas, Rose, Wildman and Dr Hannan. We acknowledge the support of University of Nottingham core facilities, including the Nottingham Biodiscovery Institute, Biosupport Unit, DeepSeq facility and Flow cytometry facility, as well as the cellular pathology department at Nottingham University Hospitals NHS Trust. We thank Prof Chris Denning for the kind donation of ReblPat human iPSCs for use in these studies. We are also grateful for the support of facilities at the John van Geest Cancer Research Centre, Nottingham Trent University and the CellGenIT team at the Wellcome Sanger Institute. WD was supported by a Clinical Research Training Fellowship (Medical Research Council MR/R017336/1) and NIHR Academic Clinical Fellowship and Academic Clinical Lectureships as well as grants from the Pathological Society of Great Britain and Ireland TSGS 1019 (WD) and CLSG 1021 (AB, MI, WD). NRFH was supported by UKRI (MRC MR/S009930/1, NC3Rs NC/X002101/1 & NC/Y000838/1 and Bowel Research UK. SCO was supported by MRC (MR/S009930/1). DS was supported by a Bowel Research UK PhD studentship. JV and SR are supported by the John and Lucille van Geest Foundation and by Nottingham Trent University’s School of Science and Technology.

## Contributions

WD, NRFH, FRAJR, RDW and MI conceived the study, contributed to experimental design and data analysis, and co-wrote the manuscript. WD performed and analysed the experiments. SM, AB and SI aided in or performed the experiments. MM and AR performed *in vivo* experiments. NH and VW performed single-cell RNA sequencing experiments. DS aided with optimising and performing immunohistochemistry. JV and SR performed NanoString Digital Spatial Profiling experiments and data analysis. AP, BC, PM, JV, SR, FS, RE and ST aided with bioinformatic analyses and interpretation of single-cell and spatial profiling studies. NRFH, SCO, SI and IT provided iPSC lines. SI, AS and IT aided in interpretation of stromal cell modulation experiments.

All authors contributed to the editing of the manuscript. The authors declare no conflicts of interest.

## Methods

### Ethical review

Animal experiments were conducted under the overview of the University of Nottingham Animal Welfare and Ethical Review Body (AWERB) and authorised by UK Home Office Project Licence numbers PPL P435A9CF8 and PP5089113.

hiPSC lines were approved and derived under University of Nottingham ethics committee number 09/H0408/74 with appropriate consent. OSTI GI4380 hiPSCs were approved for use as part of the CORGI2 trial by the Oxford/South Central Research Ethics Committee (17/SC/0079). All ethical regulations were followed throughout the study.

### Cell culture and cell lines

Human iPSCs were generated within the Nottingham Biodiscovery Institute (formerly the Centre for Biomolecular Sciences), University of Nottingham from fibroblasts harvested by punch biopsy of axillary dermis, using methods previously described (9) and were gifted to this project by Professor Chris Denning, University of Nottingham. The REBLPAT line (male) was used for most studies from passage 27 and used for experiments between passage numbers 30 and 36. BE31 and BE32 hiPSC lines (female) (34) were also used for validation. hiPSC lines were approved and derived under University of Nottingham ethics committee number 09/H0408/74. In addition, OSTI GI4380 hiPSCs (male) were used at the University of Oxford and were approved for usage as part of the Cancer Research UK CORGI2 trial by the South Central/Oxford Research Ethics Committee (17/SC/0079). Pluripotency was demonstrated by expression of pluripotency markers and lack of differentiation markers by protein/mRNA quantification.

hiPSCs were cultured in Essential 8™ medium (Life Technologies, Bleiswijk, Netherlands) to maintain pluripotency, at 37 °C, 5% CO2 in a humidified incubator. Medium was changed daily by aspirating spent medium, washing once with phosphate-buffered saline (PBS) and replacing with fresh Essential 8™ medium. Cells were cultured for three to four days to reach at least 80% confluence before passaging. In order to passage, cells were washed once in PBS, after which TrypLE Select (Life Technologies) was added for up to 5 minutes, until cleavage between cells could be seen upon microscopy, but before complete detachment from the culture surface. TrypLE Select was then quickly, but carefully aspirated, fresh Essential 8™ medium containing 10μM Y-27632 dihydrochloride (ROCK-inhibitor [ROCKi]) was added and the detached cells counted. Upon passage, approximately 30,000 iPSCs cm^-2^ were added to each culture vessel, which had been previously coated with Matrigel® at 11.1 µg cm^-2^ for at least 90 minutes incubated under standard cell culture conditions. For the first 24 hours of culture, cells were grown in medium containing 10 μM ROCKi.

Unless otherwise specified, all cell culture was performed using aseptic technique under normoxic conditions at 37 °C with 5% CO_2_ in a humidified incubator. All cells used were tested routinely for Mycoplasma at least monthly through the Nottingham Biodiscovery Institute testing programme. All media constituents and supplements are detailed in Supplementary Table 1.

### Co-differentiation to posterior endoderm and mesoderm

iPSCs were detached and passaged to an appropriate plate coated with Matrigel® (6-, 12-or 24-well; Corning) at a rate of approximately 30,000 cells cm_-2_, as described in Experimental Models. Cells were maintained for the first 24 hours in Essential 8™ medium with 10μM ROCKi to attach and proliferate as pluripotent stem cells and incubated under standard cell culture conditions. After 24 hours, the differentiation protocols 1 (novel protocol), 2 or 3 (comparator protocols) were started. Cells were assayed at various timepoints during the protocols as detailed in the results section and assay methodologies.

### MACs cell separation

Cells were detached as described above and resuspended in dissociation buffer consisting of dPBS with 1% bovine serum albumin (BSA) and 2mM ethylenediaminetetraacetic acid (EDTA). MACs separation was performed according to manufacturer’s instructions, outlined as follows. Up to 10^7^ cells were pelleted by centrifugation (300g, 8 minutes) and resuspended in dissociation buffer (100μL) with anti-EpCAM microbeads (25 μL, Miltenyi Biotec) and anti-FcR (25 μL, Miltenyi Biotec). Cells were incubated at 4 °C for 30 minutes and gently agitated every 10 minutes. The solution was then topped up to 1 mL using dissociation buffer followed by centrifugation to pellet the cells. The MACs stand and column (Miltenyi Biotec) were set up within a microbiological safety cabinet by attaching a MiniMacs™ magnetic separator (Miltenyi Biotec) to the stand and then inserting an MS column into the separator. The column was then primed with 500 μL dissociation buffer. After centrifugation, the cell pellet was resuspended in 500 μL and then added to cell column. The non-retained/flow-through cells (EpCAM-) were collected in a centrifuge tube. The column within the separator was washed 4 times with dissociation buffer (500 μL per wash). The column was then removed from the magnetic separator, 1 mL of dissociation buffer was added, and the column plunger was inserted to force retained cells (EpCAM+) into a fresh centrifuge tube. Collected cells were then washed by centrifugation twice. After the first centrifugation, if cells were being used for downstream analysis, they were resuspended in dPBS, if being used for further cell culture, cells were suspended in appropriate culture medium.

### Neutralisation of collagen

Type 1 rat-tail collagen (Life Technologies) was used at a stock concentration of 3 mg mL^-1^. To achieve a concentration of 2 mg mL^-1^ for each 1 mL of collagen gel required, 667 μL of stock was added to a tube on ice. To this, 100 μL of 10X PBS were added. To neutralise the solution, 0.025 μL of sterile 1M sodium hydroxide were added for each microlitre of collagen. The solution was topped up to 1 mL with medium (DMEM/F12), which if neutralised correctly displayed a slight pink colour of the phenol red pH indicator.

### Culture on collagen hydrogels

A method for demonstrating the growth of posterior endoderm subsequent to initial differentiation was performed on collagen hydrogels using previously published methods (21). The procedure was performed in 12-well Transwell® plates (Corning) with 0.4 μm pore size membranes. The plates were prepared by adding 500 μL of ice-cold neutralised collagen (see above) into each Transwell® insert. The plates were then incubated for 1 – 4 hours in standard culture conditions, to allow the collagen to form a semi-solid gel. Basal maturation medium consisted of DMEM / F12 (1:1) with L-glutamine (Life Technologies), 1x B27, 1x N2 (Life Technologies). This was initially supplemented with CHIR-99021 5μM, FGF-4 (Peprotech) 100 ng mL^-1^ and recombinant human Noggin 100 ng mL^-1^ (Peprotech), as well as IGF-1 (100 ng mL^-1^, Peprotech) and FGF-2 (50 ng mL^-1^, Peprotech). For the initial period of culture, all media were supplemented with ROCKi 10μM.

In the bottom of each well with a Transwell® insert 1.5mL of supplemented basal maturation medium was added. Cells were detached and re-suspended in supplemented basal maturation medium; 500,000 cells (12-well plate) were added in 500 μL of medium onto the surface of each Transwell® insert containing gelled collagen.

Cell medium was changed in both chambers every 3-4 days for the first two weeks, but with only IGF-1 and FGF-2 supplemented. Following this, the medium was aspirated from the top chamber and only replaced in the basal chamber, to create an air-liquid interface. Cultures were continued for up to 4 weeks, with media replacement in the bottom chamber every 3 – 4 days.

Following culture, medium was aspirated from the wells and replaced with 10% (*w/v*) neutral buffered formalin (NBF; Sigma Aldrich) to fix the tissues. Gels were carefully removed from the Transwell® insert and allowed to float in formalin and fixation took place overnight at room temperature, for at least 18 hours. After fixation, gels were lifted from the culture plate, blotted on tissue paper to remove excess formalin, bisected and then placed in a plastic mould. Low melting point agarose (Sigma Aldrich) was warmed to allow melting and was then laid over the gel within the plastic mould. The mould was then placed on ice to solidify the agarose. After solidifying, the agarose was trimmed and the embedded gel placed in a tissue cassette. The agarose/gel composites were fixed in NBF for a further 24 hours, before automated tissue processing (Leica) to dehydrate and take the tissues to paraffin wax. In brief, tissue cassettes were loaded onto the automated tissue processor and submerged into 50% (*v/v*) methanol in distilled water. Tissue cassettes were moved to sequentially higher concentrations of alcohol (50%, 75%, 95% and 100% [*v/v*]) for two hours each and then through three 100% xylene baths for two hours each. Finally, tissue cassettes were submerged in molten paraffin wax for two hours before being removed from the processor for embedding.

Following processing, the agarose/gel composites were embedded cut edge facing down in paraffin wax and mounted onto a cassette. Sections of cooled tissue blocks were cut using a microtome (Leica) at 3 – 4 μm, floated and collected onto poly-L-lysine coated slides. Slides were then either stored to be used later or warmed to 50 – 60 °C to melt the tissues onto the slide, before being used in histological assays described in subsequent sections.

### Stromal modulation experiments

Stromal cells were cultured on Matrigel® coated 6-well plates using standard coating parameters. The baseline media consisted of DMEM/F12 (1:1) with IGF-1 (100ng mL^-1^), FGF-2 (50ng mL^-1^) and PDGFbb (2ng mL^-1^). For TGF-β experiments, media was supplemented with TGF-β1 (1ng mL^-1^, Peprotech) or A83-01 (Sigma Aldrich, 500 nM). Cultures were maintained for up to 6 days. The same conditions were used for 3D collagen cell cultures described elsewhere but for longer periods. For purmorphamine experiments, baseline media was supplemented with purmorphamine (10 µM, Sigma Aldrich) and cultures were maintained for up to 6 days.

### RNA extraction and quantification

All RNA was extracted and purified from 6-well or 12-well plates, using a total mammalian RNA mini-prep kit (Sigma Aldrich) according to manufacturer’s instructions. Following media aspiration, wells were washed with PBS once. RNA lysis buffer containing 1% 2-mercaptoethanol was added to wells and incubated for 2 minutes at room temperature and the lysate was transferred to an Eppendorf tube, on ice. After addition of an equal volume of 70% ethanol, the mixture was transferred to an RNA binding column, and all steps for purification of RNA were performed according to the kit protocol, including an ancillary on-column DNA digestion for 15 minutes (Sigma Aldrich). Following purification, RNA was quantified using a Nanodrop™ 2000 spectrophotometer (Life Technologies).

Up to 2 μg RNA was reverse transcribed to cDNA using the Omniscript RT Kit (Qiagen, Manchester, UK) or High-Capacity Reverse Transcription Kit (ThermoFisher) according to manufacturer’s protocol. In brief, per 20 μL reaction, up to 12.5 μL of RNA template was added to 7.5 μL mastermix. The mastermix consisted of 1 μL reverse-transcriptase enzyme, 2 μL 10X RT buffer, 2 μL RT random primers, 0.8 μL 25X dNTP mix (100mM), 1 μL RNase inhibitor. All reactions were topped up to 20 μL with nuclease-free water.

All procedures were performed in a decontaminated UV-light PCR hood using nuclease free reagents and plastic consumables. Quantitative real-time PCR was carried out on the resulting cDNA. Reactions were prepared using PowerUp™ SYBR™ Green Master Mix (Applied Biosystems, Waltham, MA, USA) using custom primer pairs (Table 3; Eurofins, Ebersburg, Germany).

Primers (see supplementary table 2) were custom designed such as to span an exon-exon junctions and with a PCR product size of 70 – 150bp. All PCR sample preparation was performed in a UV-sterilised hood. Equal quantities of cDNA were added in each reaction series and at least 2 replicates per cDNA sample were performed, as well as appropriate negative controls. PCR was performed on ViiA™ 7 Real-Time PCR machine using a fast-cycling protocol consisting of an initial hold stage of two minutes at 50 °C (for activation of UNG), followed by five minutes at 95 °C (to activate hot-start Taq polymerase) followed by 40 cycles of denaturation for 1 second at 95 °C followed by annealing at an optimised temperature between 56 °C and 65 °C for 30 seconds, a separate extension step was not required. Fluorescence was read during the annealing step of each cycle. After cycling stages, a melt curve stage was included to verify the specificity of PCR amplification.

For RNA extraction and quantification performed at the University of Oxford (related to supplementary figure 2), the following procedure was used. RNeasy microkit (Qiagen, 74004) was used for RNA extraction. Extracted RNAs were incubated with DNase1 (ThermoFisher, EN0521) at 37 °C for 30 min, followed by a 10 min incubation with EDTA at 65 °C. High-Capacity cDNA Reverse Transcription Kit (Applied Biosystems, 4368814) was used to generate complementary DNA from total RNA. Quantitative real-time-PCR (qRT-PCR) was performed on LightCycler96 (Roche) with human Gapdh used as an endogenous control. The IDs of Taqman Gene expression assays (Applied Biosystems) used in this study are EpCAM (Hs00901885_m1), Cdx-2 (Hs01078080_m1), Sox17 (Hs00751752_s1), Nanog (Hs02387400_g1), Klf5 (Hs00156145_m1), Wnt2b (Hs00921614_m1), BMP4 (Hs00370078_m1), GAPDH (HS99999905-m1).

CT values for individual primers were compared with reference to a housekeeping gene using Livak’s 2^-ΔCT^ method or 2^-ΔΔCT^ method if an appropriate reference condition was available for comparison.

### Flow cytometry (EpCAM)

Cells were washed once with PBS and detached with brief TrypLE treatment. Following aspiration of the TrypLE, cells were resuspended in warm RPMI-1640 and transferred to Eppendorf tubes. The tubes were centrifuged at 300 xg for 5 minutes and then washed once with PBS, followed by further centrifugation. Reactions were performed in Eppendorf tubes and each step was followed by centrifugation at 300 xg for 5 minutes unless otherwise specified. Primary Anti-EpCAM APC conjugated antibody with isotype control was used for all experiments (Miltenyi Biotec, Surrey, United Kingdom). Cells were blocked in 3% (*w/v*) BSA in PBS for 15 minutes, followed by washing and addition of primary antibody diluted in 3% (*w/v*) BSA in PBS, with incubation for 30 minutes at 4 °C. Cells were analysed using either an FC500 or MoFlo (Beckman Coulter, Indianapolis, IN, USA) flow cytometer, with all procedures kindly optimised by the staff of the School of Life Sciences Flow Cytometry Facility, University of Nottingham (Dr David Onion and Mrs Nicola Croxall).

Data were analysed using the Kaluza Analysis software package (Beckman Coulter). Gating parameters on forward/side scatter were used consistently across all samples, and non-singlet cells were excluded from fluorescent intensity analyses to avoid overestimation. Negative and isotype controls were performed, which determined the threshold fluorescent intensity values for positive staining.

### Bulk mRNA sequencing

Cells lysates were prepared directly from plates and RNA was extracted as described earlier. After quantification by Nanodrop, samples were frozen and stored at -80 °C. Samples were diluted in nuclease-free water to obtain 2 μg total RNA in 20 μL sample volume. All samples were sent on dry ice via overnight courier to Novogene Limited for sample QC and subsequent library preparation and sequencing. Results were returned in raw FastQC format, BAM alignment files and raw and normalised count matrices for bioinformatic analysis. Subsequent analysis was performed using open-source SeqMonk and R software. Briefly, normalised gene matrices were generated using SeqMonk and initial hierarchical clustering was performed using dual approach based on intensity difference (based on log-transform) and DESeq2 packages (raw counts) to compare gene expression differences across multiple datasets. For comparison between two conditions, R was used to run DESeq2 on raw gene count matrices followed by Volcano plots based on log10 adjusted p-value and log2 fold change for individual genes. All DESeq2 analyses were performed with default parameters including correction for multiple hypothesis testing using the Benjamini-Hochberg method.

### Single-cell mRNA sequencing

Cells were dissociated as described earlier, but with a prolonged dissociation time of 8 minutes to ensure a fully dissociated single cell population. Cells were resuspended in dPBS with 1% (*w/v*) BSA to prevent intercellular adhesion. Cells were counted within the cell culture facility, using an automated counter, and resuspended at 1 million cells mL_-1_. A second count was performed within the DeepSeq facility as described below.

Single cell 3’ whole transcriptome sequencing libraries were prepared from dissociated cell suspensions using the Chromium Next GEM Single Cell 3’ Library and Gel Bead Kit v3.1, the Chromium Next GEM Chip G Single Cell Kit and the Dual Index Kit TT Set A (10X Genomics; PN-1000147, PN-1000127 and PN-1000215). Cell counts and viability estimates were obtained using the LUNA-II Automated Cell Counter (Logos Biosystems), Trypan Blue Stain, 0.4 % *(w/v)* and Luna Cell Counting Slides (Logos Biosystems; T13001 and L12001). Live cell counts were used to calculate cell input, rather than total cell count, as visual inspection of cell field on the LUNA II and the gating histogram, showed that > 90% of cells were viable and that the cell counter appeared to be counting some extracellular debris as non-viable cells. The number of input cells targeted was 3,300 cells per sample, with the aim of generating sequencing libraries from ∼ 2,000 single cells. All steps, including GEM Generation and Barcoding, Post GEM-RT Cleanup and cDNA Amplification and Library Construction were performed according to the Chromium Next GEM Single Cell 3’ Library and Gel Bead Kit v3.1 User Guide, Rev B (CG000315).

Variable steps of this protocol included using 12 cycles of cDNA amplification and 8-12 cycles of library amplification. Amplified cDNA was quantified using the Qubit Fluorometer and the Qubit dsDNA HS Assay Kit (ThermoFisher Scientific; Q32854) and fragment length profiles were assessed using the Agilent 4200 TapeStation and Agilent High Sensitivity D5000 ScreenTape Assay (Agilent; 5067-5592 and 5067-5593). Completed sequencing libraries were quantified using the Qubit Fluorometer and the Qubit dsDNA HS Assay Kit and fragment length distributions assessed using the Agilent 4200 TapeStation and the High Sensitivity D1000 ScreenTape Assay (Agilent; 5067-5584, 5067-5585).

Libraries were pooled in equimolar amounts and the final library pool was quantified using the KAPA Library Quantification Kit for Illumina Platforms (Roche; KK4824). Libraries were sequenced on the Illumina NextSeq 500 over two NextSeq 500 High Output v2.5 150 cycle kits (Illumina; 20024907) to generate > 25,000 raw reads per cell for each sample, using custom sequencing run parameters described in the 10X protocol. Following sequencing, raw outputs from the sequencer were converted to FastQC files, aligned to the reference genome (hg38) and outputted to raw and filtered count matrices using the CellRanger (v6.1.1) pipeline from 10X genomics.

Individual analysis was performed using jupyter-lab tools on Ubuntu Linux using the following packages: Scanpy (1.8.2), anndata (0.7.8), UMAP (0.5.1), NumPy (1.20.3), SciPy (1.7.3), Pandas (1.5.2), scikit (1.1.3), statsmodels (0.12.2), igraph (0.10.8), PyNNDescent (0.5.5), scvelo, cellrank, MatPlotLib, Scrublet and CellTypist (1.6.2). Briefly, samples were imported and merged. Predicted doublets were removed using Scrublet using a threshold of 0.25. Basic filtering for gene detection included presence in at least 5 cells and all cells with a minimum of 200 and maximum of 7000 genes. Genes with greater than 20% mitochondrial reads were filtered out. Cells were normalised and log-transformed and highly variable genes were then identified. Normalised expression levels next underwent principal component analysis (PCA), nearest-neighbour analysis (neighbours=15 and PCs = 10, optimised using the PCA elbow plot) and uniform mapping and approximation projection (UMAP), followed by clustering using the leiden algorithm (resolution=0.3).

Next clusters were annotated using unbiased CellTypist automated labelling. The “Developing_Human_Organs.pkl” model was selected as it is based on cells representing similar differentiation stages. Annotations were made over leiden clusters using the majority-voting method. Feature plots, dot plots and matrix plots (heatmaps) were generated using scanpy functions. The epithelial leiden cluster was extracted from the main anndata object and assessed using the “Cells_Intestinal_Tract.pkl” model within CellTypist.

### *In vivo* assays

BVA/FRAME/RSPCA/UFAW Refining Procedures for the Administration of Substances Working Group report, the NCRI Guidelines on Experimental Neoplasia, and NC3Rs Guidance for *in vivo* techniques were followed, as were the ARRIVE reporting guidelines. A total of 6 female and 12 male CD-1 NuNu mice at 6–7 weeks old were purchased from Charles River, (Margate, UK), and 6 female and 12 male Rag2^-/-^ IL2RG^-/-^ immunodeficient mice at 6-7 weeks old were purchased from Envigo, (Hillcrest), UK. Both strains were tested in order to identify if the different immune statuses of the strains had any effects on the resulting tissues.

It was essential to use only males for the testicular teratoma study, while equal numbers of males and females were used for the subcutaneous engraftment study. Numbers used per condition group were between 3-4. This was because the object of the studies was tissue generation and growth for analysis, rather than achieving statistical significance. The mice were maintained in individually ventilated cages (Tecniplast UK, Northampton, UK) within a barriered unit, illuminated by fluorescent lights set to provide a 12 h light–dark cycle (on 07.00, off 19.00), as recommended in the guidelines from the Home Office Animals (Scientific Procedures) Act 1986 (UK). The room was air-conditioned by a system designed to maintain an air temperature range of 21 ± 2 °C and a humidity of 55% ± 10%. During the study, the mice were housed in social groups, three per cage, with irradiated bedding and autoclaved nesting materials and environmental enrichment (Datesand UK, Stockport, UK). A sterile, irradiated 5V5R rodent diet (IPS Ltd., London, UK) and irradiated water (SLS, UK) were offered ad libitum. The animals’ conditions were monitored throughout the study by an experienced animal technician. After a week’s acclimatisation, the mice were initiated with cells or scaffold as detailed. At the scientific end point of the studies, the mice were killed by cervical dislocation and testes or scaffolds were removed under sterile conditions.

Animals were anaesthetised using a Ketamine (Ketaset, Animalcare, York, UK)/Medetomadine (Sedastart, Animalcare, York, UK) cocktail (Ketamine: 75mg kg^-1^ / Medetomadine: 1 mg kg^-1^) injected subcutaneously and reversed using Atipamezole (Sedastop, Animalcare, York, UK) at 1 mg kg^-1^ *s.c.*.

### *In vivo* teratoma assay

Undifferentiated iPSCs and cells differentiated according to the novel protocol and comparator protocol were dissociated and pelleted by centrifugation at 300 xg for 5 minutes. For initial experiments, either one million or two million cells were implanted to optimise conditions. Either CD1 nude or Rag2^-/-^ IL2RG^-/-^ immunosuppressed mouse models were used for optimisation. In final conditions, one million cells were implanted into Rag2^-/-^ IL2RG^-/-^ mice. Cell pellets were transported on ice to the Biosupport Unit (BSU), University of Nottingham as well as thawed Matrigel ® on ice. Immediately prior to implantation, 50 μL of Matrigel was added to the cell pellet and cells were resuspended within the matrix.

This cell suspension was then injected into the testicular subcapsular space. Mice were monitored by BSU staff throughout the period for weight and signs of ill health. Seven weeks after implantation, the mice were sacrificed and a post-mortem examination of the testicular space and abdominal cavity was undertaken. Any tumour tissue or evidence of cell growth was excised. All tissues were photographed and then placed in formalin in preparation for further histological examination.

### *In vivo* subcutaneous engraftment of populated collagen scaffolds

Tissue scaffolds were prepared from collagen following an optimised protocol in which 250 μL collagen (2mg mL^-1^) from rat’s tail was neutralised to form a hydrogel within a Transwell ® mould. After gelation, collagen gels were chemically crosslinked using a protocol adapted from Kim et al. (168). After extensive washing, a further 250 μL of neutralised collagen (2mg mL^-1^) was added to the surface of the crosslinked collagen and allowed to gel. Next, 500,000 iPSC-derived intestinal cells at day 8 were passaged to the surface of the gel in the presence of CHIR (5μM), Noggin (100ng mL^-1^), ROCKi (10μM), IGF-1 (100ng mL^-1^) and FGF2 (50ng mL^-1^) in DMEM/F12 medium supplemented with B27 and N2 supplements. The following day, medium was replaced with basal medium with additional IGF-1, FGF-2 and Rspo-1 (50ng mL^-1^). Medium was replaced again on day 4 and then on day 6 (one day before implantation). This culture medium contained IGF1 (200ng mL^-1^), FGF-2, Rspo-1 and Wnt-3a (50ng mL^-1^) to enhance to proliferation of intestinal epithelial stem cells.

On the day of transplantation, tissue grafts were carefully lifted out of the Transwell and placed in medium with the same constituents as well as ROCKi (10μM) to improve cell viability during engraftment. Tissues were engrafted by making a small incision into the flank of Rag2^-/-^ IL2RG^-/-^ immunosuppressed mice within a sterile hood. The tissue graft was inserted into the pouch formed by incision, and the incision was closed with a surgical clip. After two, three or four weeks, mice were sacrificed and the surgical site opened. Tissue from the site was removed and photographed, and fixed in formalin prior to histological examination.

### NanoString GeoMx DSP RNA profiling

Slides for NanoString analysis were prepared from formalin-fixed paraffin-embedded tissues. Multiple tissues were combined on a single slide to allow simultaneous analysis. NanoString data was collected from two separate slides consisting of representative tissues collected from different *in vivo* transplantation timepoints (slide1) and a reference normal adult human colon. Each NanoString slide consisted of two to six different pieces of tissue.

Slides were stained with pan-cytokeratin (NanoString) and DAPI stains and then hybridised with Cancer Transcriptome Atlas probes. The slides were kept hydrated at all times. The slides were then loaded onto the NanoString GeoMx data spatial profiling (DSP) instrument. Areas of illumination (AOI) were selected based on areas of relevant morphology, such as apical and basal crypt regions. Automated segmentation was also performed based on cytokeratin expression. Each AOI was selected to contain at least 100 predicted cells (nuclei) based on DAPI staining. Having selected the AOIs for each slide, automated harvesting was performed on the DSP machine and probes were collected in 96 well plates each with well-specific barcodes. Library preparation was performed according to manufacturer’s instructions and the pre-made libraries were sequenced by NovoGene Limited using Illumina NovoSeq 6000 using a PE-150 sequencing strategy on a single flow-cell lane. After sequencing, fastq files were loaded into the DSP instrument for pairing with AOIs.

To process the NanoString data, we utilised the SpatialDecon package. Prior to deconvolution, we generated cell type expression profiles from the integrated foetal gut cell atlas object, considering cells with more than 10 genes and cell types with more than 5 cells. Background counts were determined using negative probes, and the SpatialDecon algorithm was applied to estimate cell type abundances for each AOI.

Subsequently, we employed the SpatialOmicsOverlay package to visualise the data on the image data. Initially, we made necessary modifications to package functions to resolve compatibility issues. We then imported the image and annotation data along with counts into the SpatialOverlay object. AOIs were grouped based on gene expression profiles using hierarchical clustering to identify similarities, resulting in four main clusters. Then, the cell types were assigned to each AOI based on the highest cell type abundance. Since each main slide consisted of 2 to 6 different sub slides, AOIs from each main slide were annotated according to their location, such as top-right or bottom-left. To focus on individual sub slides, the SpatialOverlay object was cropped accordingly, and marker expression levels were visualised by plotting them onto the AOIs.

### Basic histology

Slides were dewaxed and rehydrated by sequential immersion in xylene, methanol and distilled water. After rehydration, slides were laid flat and tissue sections were covered with Shandon instant haematoxylin (Life Technologies) for 3 minutes. Slides were then washed in distilled water, differentiated as necessary by rapid immersion and emersion in acid-alcohol (3% HCl in 95% ethanol) and washed again in distilled water. Submersion in Scott’s tap water (1% w/v magnesium sulphate and 0.067% w/v sodium bicarbonate in distilled water) was performed for 5 minutes to blue the haematoxylin, followed by further washing in distilled water. The slides were stained with Shandon instant eosin (Life Technologies) for 30 seconds, briefly washed in distilled water and then dehydrated by sequential immersion in ethanol and xylene. Slides were mounted with DPX and a coverslip was placed over the tissue.

For Alcian blue staining, tissues were similarly rehydrated. Slides were incubated in 3% (*v/v*) acetic acid solution for 1 minute to acidify the tissue and then immersed in Alcian blue solution for 15 minutes. After washing in distilled water, slides were differentiated as necessary in acid-alcohol, washed in distilled water and then tissues were covered with nuclear fast red solution (Sigma Aldrich) for 5 minutes for counterstaining. Following washing, slides were dehydrated and mounted as described above.

### Immunohistochemistry

Slides were dewaxed and rehydrated as described above. Owing to previous fixation, antigen retrieval was performed by incubating slides in simmering sodium citrate buffer (pH6) for 20 minutes; a microwave set on low power was used to maintain temperature (approximately 95 °C). Once cooled, slides were mounted under water onto Shandon Sequenza® coverplates (Life Technologies) and placed into a Sequenza® rack. Slides were washed three times with 200 μL TBST (Tris-buffered saline with 0.01% Tween-20).

All immunohistochemistry was performed using the Novolink™ polymer detection system according to manufacturer’s instructions; all volumes used were 100 μL and washing was performed with TBST. Antibodies and conditions are detailed in Supplementary Table 3. Briefly, following protein and peroxidase blocking steps and washing, 100 μL of diluted antibody solution (in TBST) was added per slide; see Table 4 for details of antibodies used and concentrations. Following overnight incubation in the primary antibody solution at 4 °C, slides were washed and exposed to the post-primary solution for 1 hour at RT, polymer-peroxidase for 30 minutes, 3’,3’-diaminobenzidine (DAB) for 5 minutes and modified haematoxylin for 5 minutes. After this, slides were dehydrated and mounted as described above. Some IHC was performed by the Nottingham University Hospitals Cellular Pathology Department, according to standard operating procedures, which are available upon request; any such assays are indicated within the text.

### mRNA *in situ* hybridization

To detect expression of stem cell marker *LGR5* in the cultured cells on day-08 of the differentiation, EpCAM^+^ cells were fixed in 4% paraformaldehyde for 30 min at RT on a glass slides and dehydrated through ascending concentrations of ethanol (50%, 70%, 100%) and stored at -20C. On the day of the staining, the cells were rehydrated by submerging the slides in 70% ethanol (2 min), 50% ethanol (2 min) and 1X PBS (10 min) at RT. The cells were then perforated by rinsing the slides in 1X PBST (1X PBS containing 0.1% Tween20) for 10 min at RT. After washing in 1X PBS (5 min), cells were incubated for 10 min at RT in RNAscope hydrogen peroxide solution (Cat # 322335). Then, slides were washed in 1X PBS and incubated in RNAscope Protease III solution, diluted 1:15 in 1X PBS, for 10 min at RT inside a humidity control chamber. After washing the slides in 1X PBS again, the manufacturer’s protocol for RNAscope Multiplex Fluorescent Kit v2 (Cat # 323110) was followed starting from the probe hybridisation step. *LGR5* mRNA signal was detected by using RNAscope catalogue probe 584631.

To detect *LGR5* mRNA expression in formalin-fixed paraffin-embedded organoids, 4 um thin sections were stained with RNAscope *LGR5* probe (Cat# 584631), following the manufacturer’s guideline, using RNAscope Multiplex Fluorescent Kit v2 (Cat # 323110). For the co-detection of *LGR5* mRNA, EPCAM and KI67 proteins, an immunofluorescence staining was performed after mRNA ISH following the protocols described by Montazid et al. (https://www.nature.com/articles/s41467-023-44138-6). Concentrations of antibodies used in this protocol were: 1:500 for EpCAM (Abcam, ab71916) and 1:500 for Ki67 (Cell Signaling, 12202).

### Statistical analysis

Statistical analyses were performed if appropriate to determine statistically significant differences. The student’s t-test (unpaired, two-tailed) or one-way ANOVA tests were used for continuous data. Unless otherwise specified, the following symbols are used to indicate statistical significance: * p < 0.05, ** p < 0.01, *** p < 0.001, **** p < 0.0001. Numbers of replicates, statistical tests and summary statistics are indicated in the methods and results sections or figure legends.

All statistical analyses and graphs were performed and produced using the following software packages: Microsoft Office 365 Excel, GraphPad Prism 9, R x86 v4.0.3/4.1.1/4.3 and Jupyter-lab v3.3.0. Image analysis was performed using Leica LASX, ImageJ/Fiji, QuPath and Adobe Photoshop.

## Data availability

All raw and processed sequencing data will be made available on EMBL Array Express upon acceptance for publication.

## Code availability

All code is available at http://github.com/dalleywater/NUGut.

---

**Suppl. Table 1:**

**Culture medium and supplements**

List of supplements is available upon request to the corresponding author.

**Suppl. Table 2:**
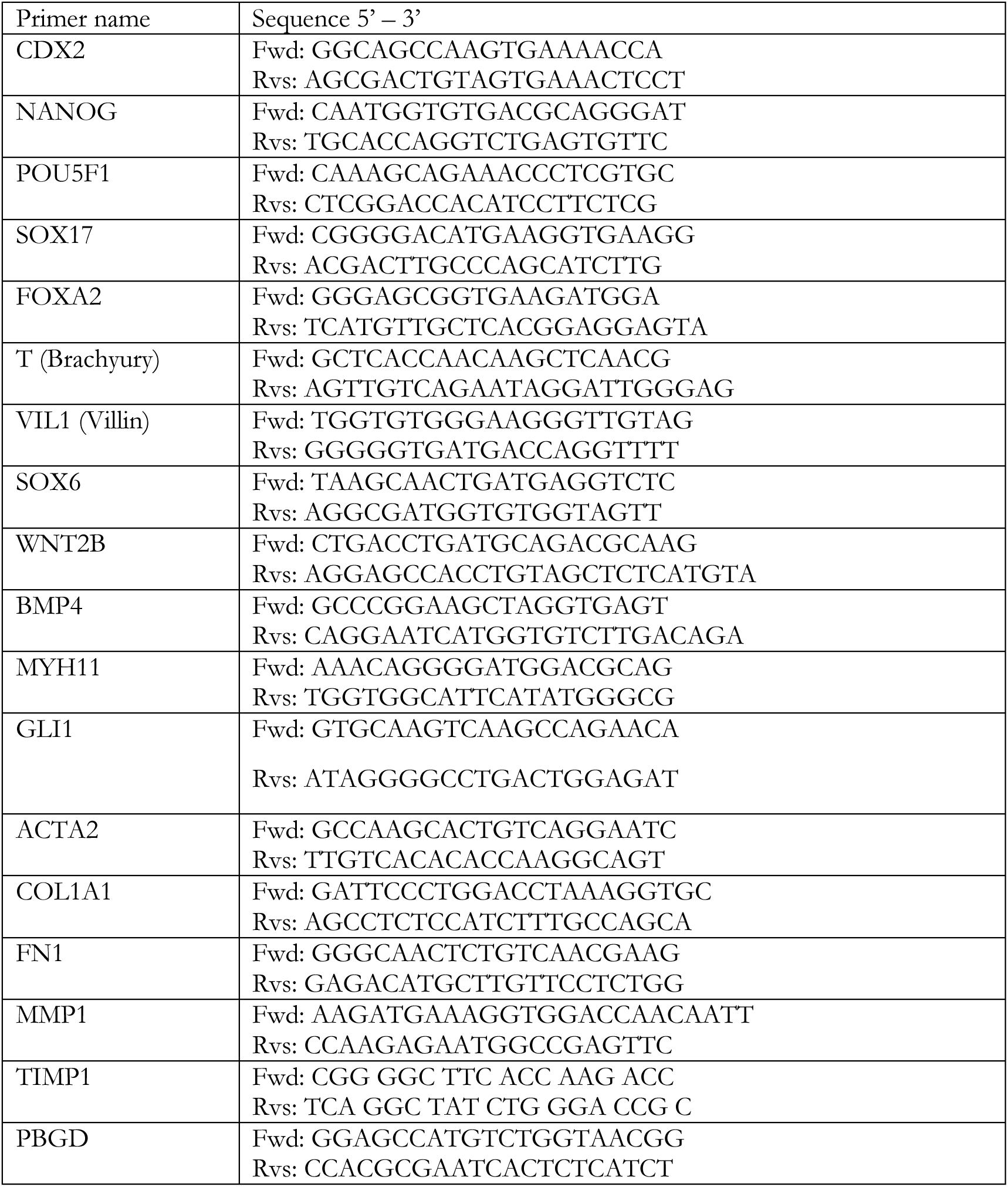
qRT-PCR primers.

**Suppl. Table 3:**
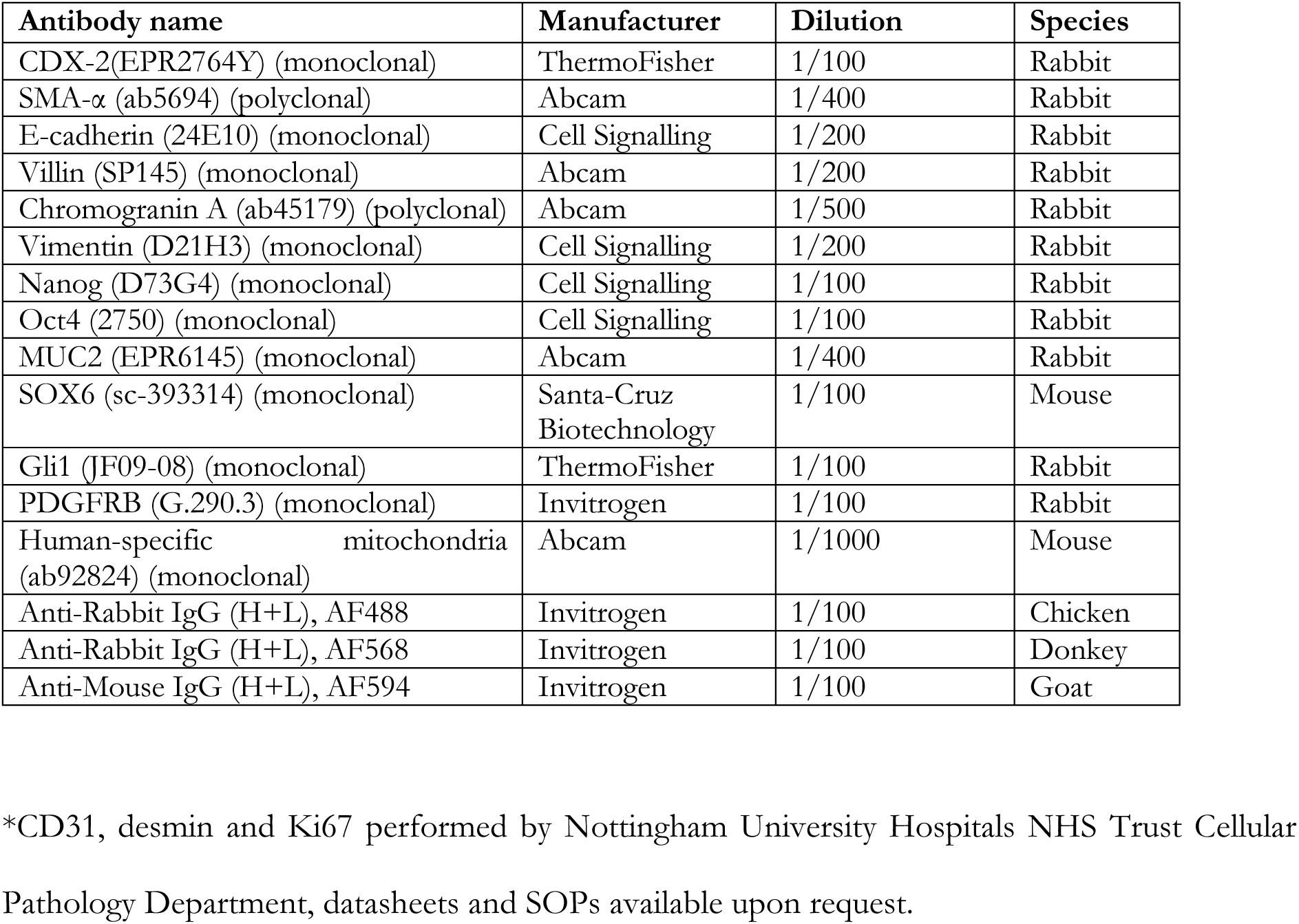
Antibodies.

## Notes

### Competing Interest Statement

The authors have declared no competing interest.

### Summary of Updates

Figures reorganised to improve clarity of data. Update to single cell annotation using latest CellTypist model (Fig. 3).

